# Data indicate the importance of expert agencies in conservation policy

**DOI:** 10.1101/424911

**Authors:** Michael J. Evans, Jacob W. Malcom, Ya-Wei Li

## Abstract

Data on the implementation of laws and policies are essential to the evaluation and improvement of governance. For conservation laws like the U.S. Endangered Species Act (ESA), such data can inform actions that may determine the persistence or extinction of species. A central but controversial part of the ESA is section 7, which requires federal agencies to conserve threatened and endangered species. One way they do this is by consulting with expert agencies for the ESA, the U.S. Fish and Wildlife Service (FWS) and the National Marine Fisheries Service (NMFS), on actions they may undertake that impact listed species. Using data from all 24,893 consultations recorded by NMFS from 2000 through 2017, we show that federal agencies misestimated the effects of their actions on listed species in 21% of consultations, relative to the conclusions reached by NMFS. In 71% of these cases the federal agency underestimated the effects of their action. Those discrepancies were particularly important for the conservation of 14 species in 22 consultations, where the agency concluded that its action would not harm a species, while NMFS determined the action would jeopardize the species’ existence. Patterns of misestimation varied among federal agencies, and some of the agencies most frequently involved in consultation also frequently misestimated their effects. Jeopardy conclusions were very rare—about 0.3% of consultations—with a few project types more likely to lead to jeopardy. These data highlight the importance of consultation with the expert agencies and reveal opportunities to make the consultation process more effective.

**SIGNIFICANCE STATEMENT:** The US Endangered Species Act is the strongest environmental law any nation has enacted to conserve imperiled species. However, policy debates over how the Act should be implemented continue to this day. This study provides the first comprehensive evaluation of how the National Marine Fisheries Service (NMFS) implements one of the Act’s most important conservation programs – consultations under section 7. Our results reveal novel insights into the importance of NMFS role in ensuring federal actions do not jeopardize the existence of listed species. By using data to inform policy debate, we identify approaches to implementing section 7 that would undermine the conservation of imperiled species, and those that could improve the efficiency of the program without sacrificing these protections.

## INTRODUCTION

Data-driven decision-making is an important and growing theme in society. In governance, data can provide an accurate picture of how laws and policies are implemented, highlight real successes, and identify shortcomings that need to be addressed^1,2^. The role of regulation in society is often framed as a choice between polarized extremes, and debates about regulation can become binary and ideological^3^. Data can temper extreme rhetoric and false dichotomies, providing a more nuanced evaluation of the benefits and drawbacks of regulatory approaches^4^, offer evidence for competing alternatives, and identify areas for compromise^5^. There is a pressing need to use data to inform environmental policy and implementation because both often involve opposing ideals.

The strongest legislation any nation has enacted to conserve imperiled species is the U.S. Endangered Species Act (ESA). A key source of this strength is section 7 of the ESA, which requires that federal agencies consult with the U.S. Fish and Wildlife Service (FWS) or National Marine Fisheries Service (NMFS; together “the Services”) to ensure that actions the agencies take, fund, or permit will not jeopardize the existence of any species on the endangered species list or adversely modify these species’ critical habitat (Box 1). Both listed species and any Distinct Population Segments (DPS) or Evolutionarily Significant Units (ESU) - distinct segments of a species that can be independently listed under the ESA^6^ - require consultation under section 7^7^. The consultation requirement is important for the conservation of imperiled species and is the primary regulatory protection in the ESA for plants^8^. At the same time, section 7 is criticized as inefficient and burdensome by some parties, although recent research using consultation data from FWS indicates this criticism is often overstated^9^. These conflicting views raise an important question: How can the ESA be most effective at conservation and cost-efficient to implement?

### Box 1: The Section 7 Consultation Process

Section 7 consultation typically begins with informal consultation after a federal agency determines that a proposed action may affect a listed species. If informal consultation is initiated by an agency, FWS or NMFS reviews whether the proposed action is “likely to adversely affect” (LAA) a species. If the Services determine that the proposed action is “not likely to adversely affect” (NLAA) a species or critical habitat, consultation can end at this stage. If an LAA determination is made for a species or critical habitat, “formal consultation” is initiated, during which the Services evaluate whether the proposed action will violate the ESA prohibitions on jeopardizing species (i.e. appreciably reducing a species’ probability of survival) or destroying/adversely modifying their critical habitat. If either of these outcomes is likely, the Service must suggest “reasonable and prudent alternatives” that can be implemented by the federal agency to reduce or offset harm caused by the proposed action. If no alternatives are available, the federal agency cannot proceed with the action without violating the ESA unless it obtains an exemption from the Cabinet-level Endangered Species Act Committee. We hereafter refer to federal agencies engaged in the consultation process as “action agencies,” and the actions requiring consultation as “proposed actions.” Consultations may cover multiple species, each of which may be affected differently by a proposed action. We refer to the individual species effects of an action as a “determination.” Thus, many consultations have multiple determinations.

For decades, government agencies, politicians, and the public have offered competing approaches to improve implementation of the ESA^10,11^. One common type of proposal kis to reduce the role that FWS and NMFS play in evaluating the effects of proposed federal actions by allowing action agencies to make that determination for itself. These “self-consultation” approaches have included the 2004 pesticide counterpart rule^12^, the National Forest counterpart rule^13^, and the alternative consultation regulations during the G.W. Bush administration^14^. Interest in reducing expert agency involvment continues to this day: exemptions from section 7 consultation for forest management and pesticide registrations were proposed in a U.S. House of Representatives draft of the 2018 Farm Bill^15^, and the 2018 revisions to regulations on interagency cooperation proposed by the Departments of the Interior and Commerce include an “Expedited Consultation” section^16^. A critical, outstanding question is whether these alternatives effectively conserve species protected by the ESA or simply alleviate conservation obligations and streamline compliance. This question has never been quantitatively evaluated despite the controversy surrounding this issue, because the data have not previously been available.

Here we provide the first data-driven examination of the modern consultation program of NMFS, the expert agency responsible for evaluating federal actions occurring in marine environments or affecting most anadromous fishes. Our goal was to answer two fundamental questions about how consultations occur. First, how often do federal agencies misestimate the effects of their actions on threatened and endangered species? This is important to know for assessing whether ideas such as self-consultation will maintain or reduce protections for endangered and threatened species. Second, what are the general patterns of consultation outcomes, including jeopardy and adverse modification determinations? This is critical for understanding the conservation impacts of the consultation program and predicting the effects of proposed changes.

We first show that over the past 17 years, more than one-fifth of consultations have included proposed determinations from action agencies that could result in the under- or over-protection of species relative to the expert NMFS determinations. Certain agencies and types of actions are more likely to miss the mark, but the results strongly indicate that in general, expert agency review can be critical for protecting threatened and endangered species while self-consultation could compromise this purpose of the ESA. Second, NMFS rarely determined that federal actions would jeopardize species or adversely modify critical habitat, and no actions were stopped as a result of NMFS finding jeopardy or adverse modification without reasonable and prudent alternatives. Together with quantitative and qualitative descriptions of consultations, such as patterns of jeopardy determinations, the data point to strengths of the section 7 program and can be used to identify potential improvements in its implementation.

## RESULTS

The Public Consultation Tracking System (PCTS) database shows that NMFS biologists recorded 19,826 informal and 4,934 formal consultations (19.9% formal) from January 2000 through June 2017. These numbers exclude consultations recorded as technical assistance over the same period. While the number of informal consultations increased (*Δper year* = 28.71, *SE* = 7.21, *F_16,1_* = 15.84, *P* = 0.001), the number of formal consultations has remained relatively constant (*Δper year* = −0.53, *SE* = 2.46, *F_0,1_* = 0.05, *P* = 0.833) over time. The number of formal and informal consultations differed geographically by NMFS region (*X^2^_5_* = 11,438.8, *P* < 0.001), and were highest within the West Coast region (3,589 formal, 11,664 informal). Consultations were unevenly distributed among species (*X^2^_9_* = 11,872.6, *P* < 0.001), federal agencies (*X^2^_9_* = 69,853.0, *P* = < 0.001), and work types (*X^2^_9_* = 19,185.7, *P* < 0.001). The species most commonly consulted on (Fig. 1A) were chinook salmon (*Oncorhynchus tshawytscha*) and steelhead trout (*Oncorhynchus mykiss*). The most frequently consulting agency was the Army Corps of Engineers, with a consultation rate ~ 6 times higher than the next-closest agency, the Forest Service (Fig. 1B). The most common work type requiring consultation was ‘waterway’ (Fig. 1C), which includes activities like flood control, bank stabilization, dredging, and dock construction.

**Figure 1.**
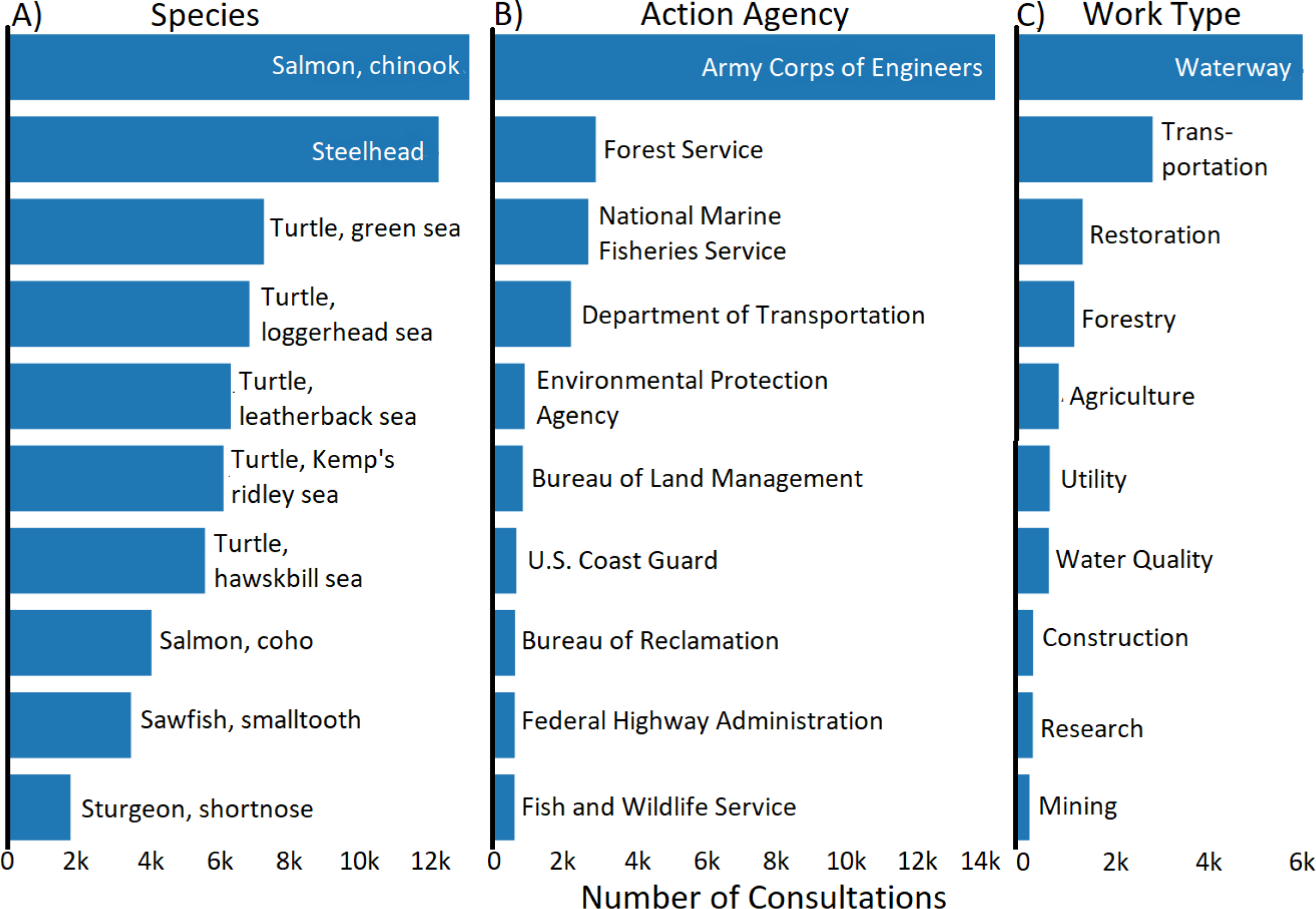
Consultation frequencies differ substantially between species (A), federal agencies (B), and types of activity (C). Frequencies of section 7 consultations conducted by the National Marine Fisheries Service involving different species, action agencies, and work types between 2000 and 2017. The ten most frequent members of each group are shown.

To evaluate how often action agencies’ determinations aligned with NMFS species experts’ determinations, we compared agencies’ proposed determination to the NMFS final determinations. In cases of disagreement, we assume the NMFS analysis to be more accurate because NMFS is the expert wildlife agency, although this assumption may not be true in all cases. NMFS disagreed with the proposed determinations of action agencies less frequently (21% of the time) than they agreed (Table 1). Among these discrepancies, action agencies more frequently (71%) underestimated the effects of proposed actions than they overestimated effects (Fig. 2). The most frequent form of discrepancy occurred when an action agency proposed an NLAA determination and NMFS subsequently made an LAA determination, leading to either a jeopardy or a no jeopardy determination (Table 1). Agencies differed in the patterns of disagreement with NMFS on the effects of proposed actions (Fig. 2). Among action agencies with at least 20 determinations, the Environmental Protection Agency (*D_122_* = 0.37, *P* < 0.001) and National Park Service (*D_39_* = 0.16, *P* = 0.005) tended to underestimate the effects of proposed actions. Conversely, the Army Corps of Engineers (*D_2508_* = 0.04, *P <* 0.001) and Federal Energy Regulatory Commission (*D_83_* = 0.13, *P <* 0.001) tended to overestimate effects. The Federal Emergency Management Agency (*D_52_* = 0.12, *P* = 0.028) both over and underestimated effects. The Bureau of Land Management (*D_195_* = 0.07, *P* = 0.016), Forest Service (*D_469_* = 0.07, *P <* 0.001), and NMFS (*D_1525_* = 0.05, *P <* 0.001) were in agreement with NMFS more often than were other agencies on average (Fig. 2).

**Table 1.**
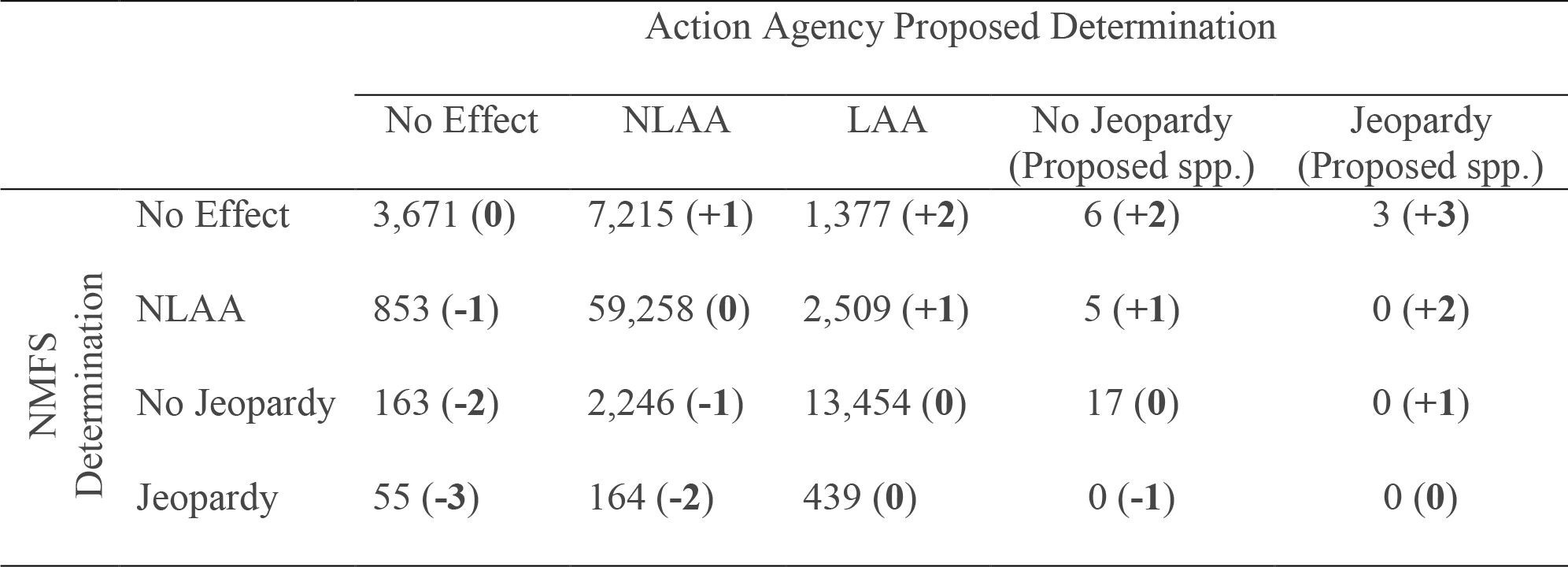
Frequencies of determinations proposed by action agencies vs. final determinations made by NMFS during section 7 consultation from 2000 to 2017. Bold numbers show the ‘discrepancy’ score assigned to a given combination, indicating the degree of agreement (positive values) or disagreement (negative values).

|  |  | Action Agency Proposed Determination |  |  |  |  |
| --- | --- | --- | --- | --- | --- | --- |
|  |  | No Effect | NLAA | LAA | No Jeopardy<br>(Proposed spp.) | Jeopardy<br>(Proposed spp.) |
| NMFS<br>Determination | No Effect | 3,671 (**0**) | 7,215 (+**1**) | 1,377 (+**2**) | 6 (+**2**) | 3 (+**3**) |
|  | NLAA | 853 (**-1**) | 59,258 (**0**) | 2,509 (+**1**) | 5 (+**1**) | 0 (+**2**) |
|  | No Jeopardy | 163 (**-2**) | 2,246 (**-1**) | 13,454 (**0**) | 17 (**0**) | 0 (+**1**) |
|  | Jeopardy | 55 (**-3**) | 164 (**-2**) | 439 (**0**) | 0 (**-1**) | 0 (**0**) |

**Figure 2.**
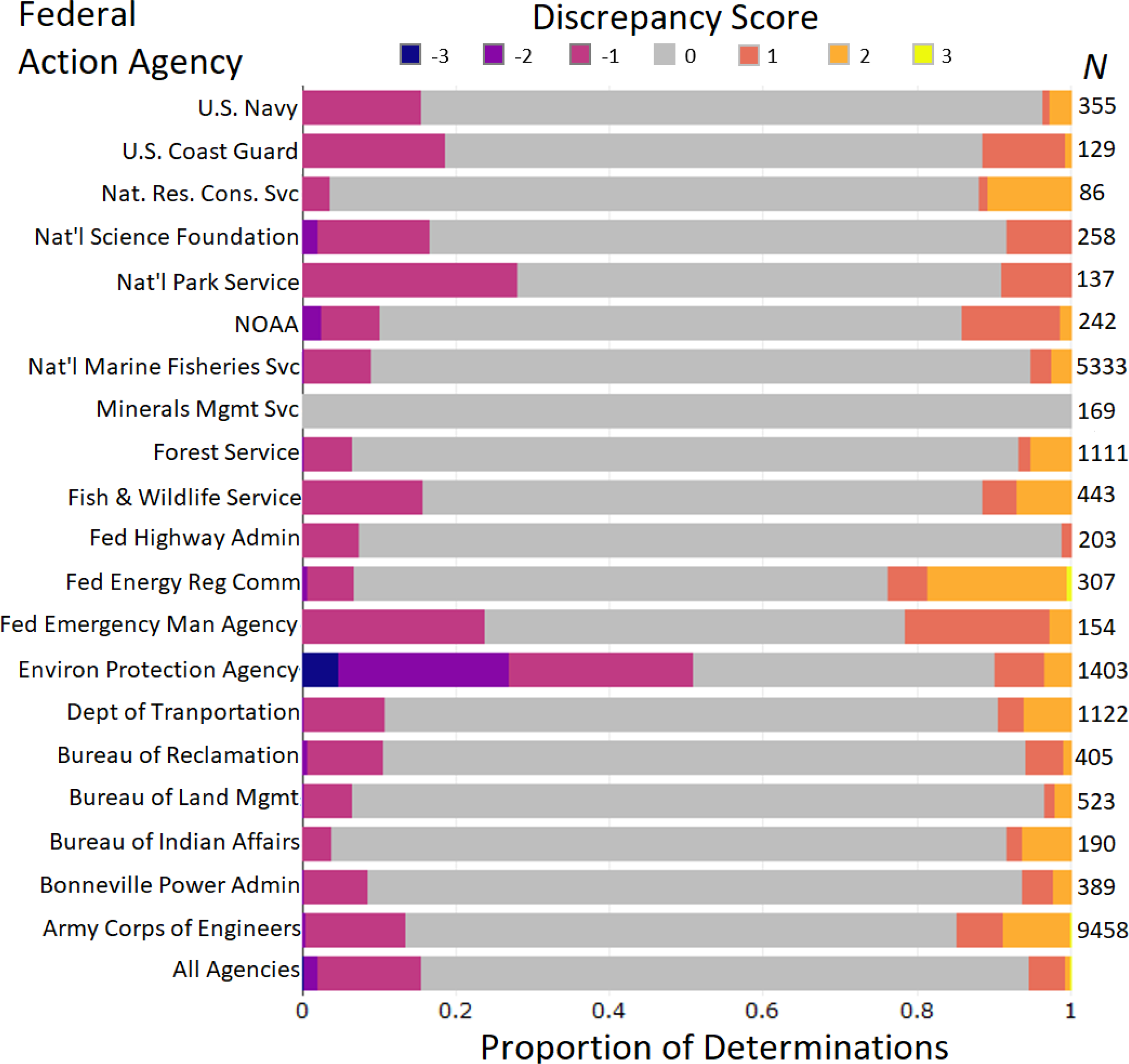
Federal agencies vary in rates of disagreement with NMFS during section 7 consultation. Ordinal ‘discrepancy’ scores indicate the degree of disagreement on determinations between action agencies and NMFS during section 7 consultations between 2000 and 2017. Positive values indicate overestimation of effects by an action agency, and negative values indicate underestimation. Bar length represents the percentage of determinations by an agency receiving each score, and ‘N’ provides the number of determinations.

To understand the causes and consequences of misestimated determinations that ultimately concluded with jeopardy, we explored jeopardy and adverse modification conclusions in greater detail. Of the 4,934 formal consultations, 72 (1.5% of formal and 0.3% of all consultations) resulted in jeopardy findings and 55 (1.1% of formal, 0.2% of all consultations) resulted in findings of adverse modification of critical habitat. These consultations consisted of 641 jeopardy and 503 adverse modification determinations. Three consultations resulted in adverse modification without jeopardy and 37 resulted in jeopardy without adverse modification. All projects could proceed if the permittee adopted reasonable and prudent alternatives to minimize or partially offset the adverse effects of the project. The rate of consultations that ended in jeopardy was constant over time (*Δper year* = −0.001, *SE* = 0.001, *F_1,15_* = 4, *P* = 0.640). Rates of jeopardy determinations differed among species (*X^2^_9_* = 16.15, *P* = 0.064), and rates of jeopardy consultations differed among work categories (*X^2^_8_* = 153.69, *P* < 0.001). Federal actions related to fisheries management and pest control were more likely to result in jeopardy than other work types (Fig. S1A). Among species with at least 10 consultations, the Cook Inlet DPS of beluga whale (*Delphinapterus leucas*) had the highest rates of jeopardy determinations (Fig. S1B), although Pacific salmonid species had the greatest number of jeopardy determinations (Table 2).

**Table 2.** The ten species with the greatest number of jeopardy determinations in formal NMFS section 7 consultations between 2000 and 2017. The number of distinct DPS/ESUs involved in jeopardy determinations is shown parenthetically.

| Common Name | Jeopardy Determinations | All Determinations | % |
| --- | --- | --- | --- |
| Green sea turtle (1/5 DPS) | 9 | 778 | 1.2 |
| Kemp’s ridley sea turtle | 9 | 614 | 1.5 |
| Killer whale (1/1 DPS) | 9 | 186 | 4.8 |
| Loggerhead sea turtle (2/8 DPS) | 10 | 768 | 1.3 |
| North Atlantic right whale | 10 | 176 | 5.7 |
| Leatherback sea turtle | 11 | 703 | 1.6 |
| Chum salmon (2/4 ESU) | 16 | 564 | 2.8 |
| Sockeye salmon (2/7 ESU) | 24 | 675 | 3.6 |
| Coho salmon (5/7 ESU) | 31 | 1490 | 2.1 |
| Steelhead (13/15 DPS) | 60 | 3012 | 2.0 |

To evaluate whether particular types of actions led to jeopardy determinations, we tested for over- or under-representation of species-work type combinations among jeopardy determinations. There was a disproportionately high number of jeopardy determinations resulting from proposed actions categorized as “agriculture” affecting chinook salmon (*Effect* = 8.6, *P* = 0.003), coho salmon (*Effect* = 10.9, *P* < 0.001), and steelhead (*Effect* = 8.7, *P* < 0.001; Fig. 3). The agriculture work type includes pesticide registration, irrigation, and grazing allotment decisions. Blue (*Balaenoptera musculus*), humpback (*Megaptera novaeangliae*), fin (*B. physalus*), North Atlantic right (*Eubalaena glacialis*), sei (*B. borealis*), and sperm whales (*Physeter macrocephalus*), as well as leatherback sea turtles (*Dermochelys coriacea*), were disproportionately jeopardized by actions in the “fishery” category (*Effect* > 1.5, *P* < 0.05; Fig 3). Finally, ringed seal (*Phoca hispida*) were disproportionately jeopardized by actions categorized as “utility” (*Effect* = 1.0, *P* = 0.038; Fig. 3), which includes hydropower, pipeline, and transmission line construction and maintenance, and gulf sturgeon (*A. oxyrinchus desotoi*) by actions categorized as “ocean” (*Effect* = 1.0, *P* = 0.030), which includes shoreline stabilization, geotechnical exploration, and waste disposal. Both coho salmon and steelhead were jeopardized less than expected by fishery actions (*Effect <* −4.5, *P <* 0.007), and both chinook and coho salmon less than expected by waterway actions (*Effect <* −4.0, *P* <0.05; Fig. 3).

**Figure 3.**
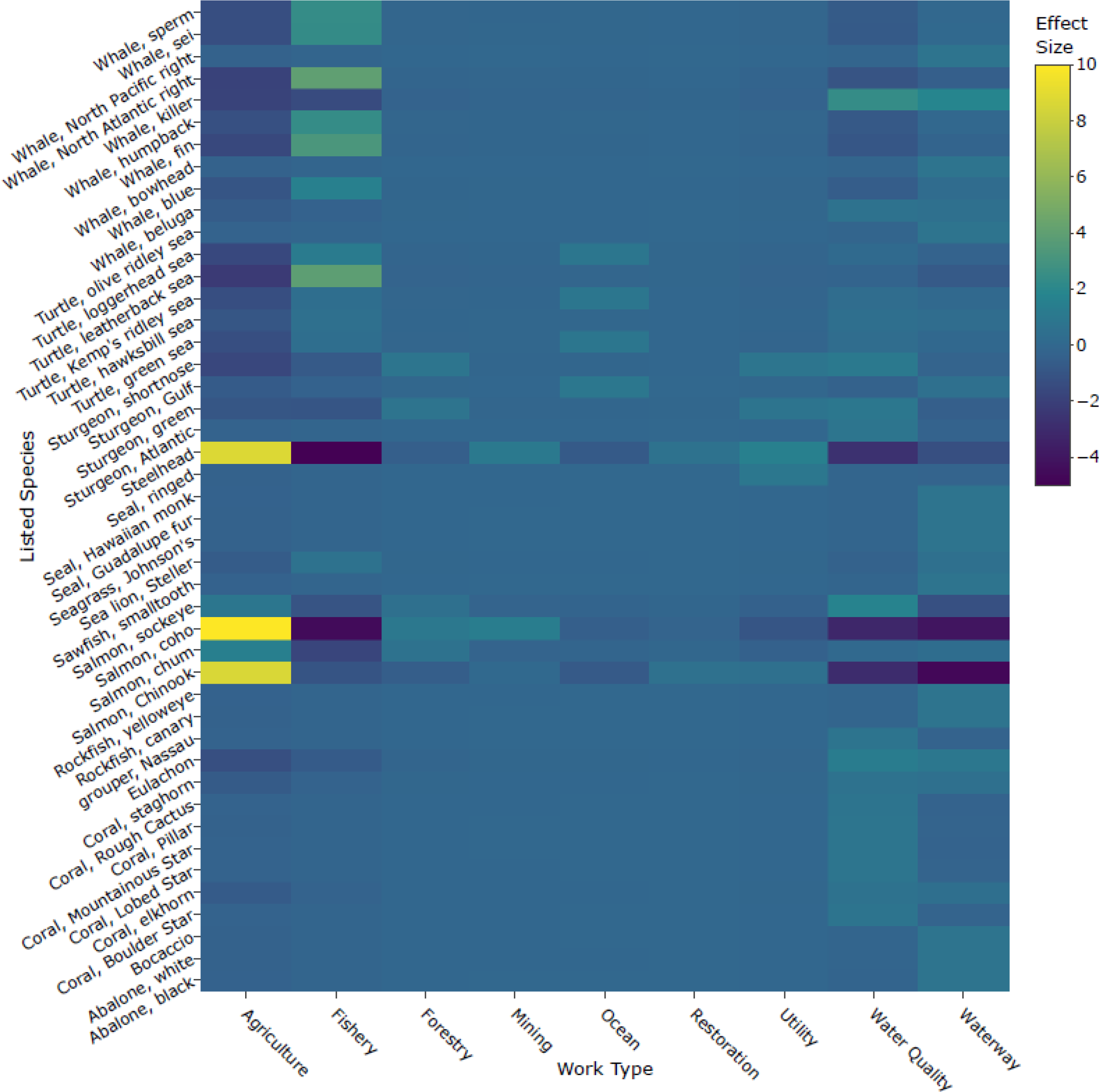
Different species are vulnerable to specific types of actions. Heatmap displays differences between the observed and expected frequency of jeopardy determinations for combinations of species and work types during section 7 consultation between 2000 to 2017. Higher values indicate combinations for which jeopardy determinations occurred more frequently than would be expected under a random association.

Chinook salmon and steelhead had the highest probability (0.42) of being jeopardized by the same action (“co-jeopardization”) among all species (Fig. 4), and all Pacific salmonids exhibited significant co-jeopardization (*Effect* > 2, *P* < 0.024). Additionally, the Southern Resident DPS of killer whale (*Orcinus orca*) were co-jeopardized with all Pacific salmonids (*Effect* > 4.4, *P* < 0.018), except coho salmon. Finally, significant co-jeopardization occurred between green sturgeon (*Acipenser medirostris*) and sockeye salmon (*Effect* = 2.9, *P* = 0.009), and eulachon (*Thaleichthys pacificus*) and coho salmon (*Effect* = 2.4, *P* = 0.024; Fig. 4).

**Figure 4.**
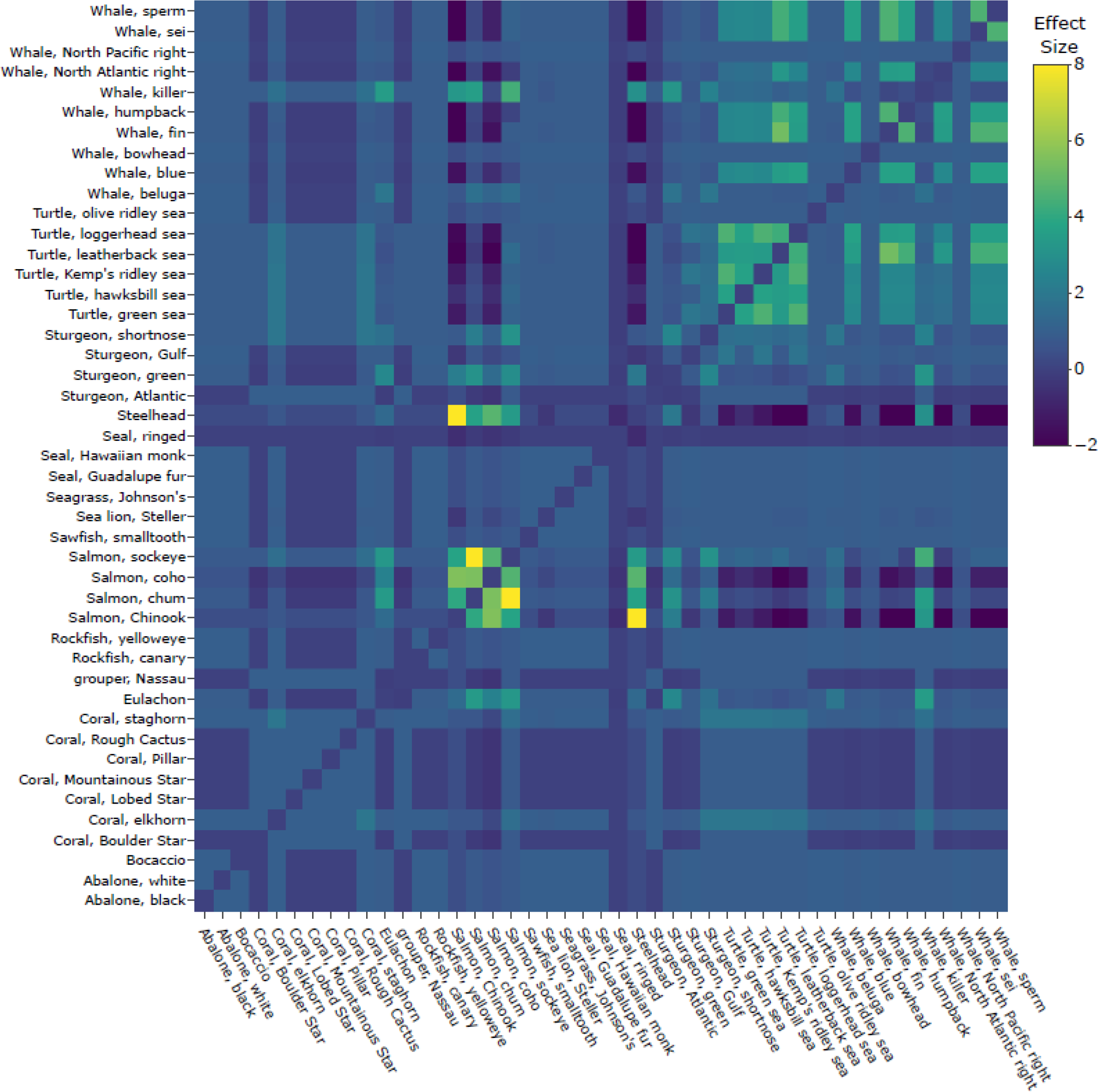
Some pairs of species are likely to be jeopardized by the same action. Heatmap displays differences between the observed and expected frequency with which jeopardy determinations were made for pairs of species during the same NMFS section 7 consultation between 2000 to 2017. Higher values indicate species pairs for which observed co-jeopardizations were greater than would be expected under a random association.

## DISCUSSION

Using data to critically evaluate the efficacy and efficiency of laws and regulations can help clarify contentious topics and guide the development of future policy. Consultation between federal agencies and species experts at the Services is one of the most important provisions in the ESA, and often the most controversial^17,18^. Our analysis of NMFS consultation data provides the first quantitative evidence of the importance of having species experts evaluate the potential effects of proposed federal actions, rather than relying solely on action agency staff. The data illustrate that even though the vast majority (99.7%) of federal actions proceed without substantial changes, rare instances of jeopardy determinations made by species experts at NMFS were critical to ensuring that federal agencies did not authorize actions that would harm or even jeopardize listed species. The results indicate that recommendations to reduce the role of expert agencies in the consultation process may compromise the conservation of imperiled species, and also point to possible approaches to improve the efficiency of consultation without sacrificing species protections.

Our results show that excluding the species experts at NMFS from the consultation process could have been detrimental to the conservation of certain threatened and endangered species. The purpose of the ESA is to conserve imperiled species, and section 7 consultations are the primary mechanism through which the ESA ensures that federal agencies do not compromise this purpose^19^. The data show that allowing self-consultation by action agencies would have resulted in action agencies failing to recognize that their proposed actions would likely jeopardize a listed species or destroy or adversely modify critical habitat, violating (albeit perhaps unintentionally) their duties to ensure against such risks. While NMFS agreed with the majority of action agencies’ proposed determinations, agreement rates varied substantially depending on the action agency, type of action, and species. The potential impact of these erroneous consultation outcomes is severe: if action agencies had been allowed to self-consult, actions resulting in almost one-third (219) of the jeopardy determinations made by NMFS would not have received thorough biological analysis because the action agencies had made a no effect or NLAA determination. Without NMFS involvement, these instances would have authorized 22 actions that jeopardized 14 species, many of which are economically important (e.g., commercially harvested salmonids).

The details of cases of disagreement between action agencies and NMFS offer insights into the implications for future policy. These cases were primarily limited to a subset of species and work types in which wide-ranging actions had the potential to adversely affect multiple, spatially-overlapping species. The federal registration of pesticides (categorized under the agriculture work type) made up a disproportionate percentage of jeopardy findings for multiple species of Pacific salmonids, and the authorization of Atlantic fisheries plans made up a disproportionate percentage of jeopardy findings for several whale and turtle species. The spatial extent of these proposed actions and the spatial overlap of affected species meant that such actions would have potentially severe consequences if their effects were underestimated. This emphasizes the importance of involving species experts in the consultation process. Most federal agencies do not have the same biological expertise or dedicated resources to conduct an equally thorough and informed evaluation of the conservation impacts of their actions as does NMFS. Furthermore, it is likely that agencies whose priority is not the protection of imperiled species may be motivated to expedite projects that fulfill their institutional mission. Therefore, checks for potentially harmful federal actions–like those currently provided by expert evaluation–are crucial for the ESA to prevent the extinction of species.

Patterns of concurrence between action agencies and NMFS can also be used to identify and inform opportunities for increased efficiencies in the section 7 consultation process that do not sacrifice species protections. Because an LAA finding triggers the expenditure of additional effort for formal consultation, providing clear guidance at this stage might reduce the consultation workload considerably without undermining conservation. The LAA determination was the most common type of discrepancy between NMFS and action agencies, accounting for 53% of misestimates, and presents an opportunity to improve consultation efficiency. The best example of this was the case of waterway activities. These actions were most commonly initiated by the Army Corps of Engineers and Federal Emergency Management Agency, which often overestimated the effects of proposed actions. Such overestimation resulted in 159 unnecessary formal consultations because NMFS ultimately concluded that the species were not affected or were unlikely to be adversely affected. This is an example where explicit standards for LAA thresholds and/or programmatic consultation could reduce instances of disagreement and conserve agency resources. Policy guidance to address the current lack of detailed, quantitative standards for the LAA threshold could be very fruitful. The interim approaches for pesticide assessment^20^ and the consultation keys for woodstork^21^ provide two different models of how the Services have clarified the LAA threshold.

Patterns of jeopardy determinations among species-work type combinations provide evidence for the benefit of advanced planning to improve both species outcomes and consultation efficiency. The best example comes from actions related to fisheries management, which resulted in fewer jeopardy determinations than expected for multiple Pacific salmonid species. On the surface, this result was surprising, because we expect these types of actions to negatively affect anadromous species. However, in the case of fishery management, a rule issued under section 4(d) of the ESA allows commercial use of some listed salmonid species^22^ and sets quantitative standards for developing and approving fishery management plans for these species. This process front-loaded much of the analysis of effects for similar actions, expediting subsequent consultation and reducing the probability that proposed actions would jeopardize the species. There are likely other, similar opportunities to improve how other sections of the ESA are implemented.

The NMFS data show that jeopardy and adverse modification determinations are very rare, and we know of no instance in which such a determination stopped a project because alternatives were unavailable. We note, however, that the very low rates of jeopardy and adverse modification from NMFS (<2% of formal consultations) are higher than those from FWS (<0.1% of formal consultations)^7^. One possible explanation is that DPS/ESUs are more common among species managed by NMFS than FWS, and the effects of proposed actions may be more likely to cross a jeopardy threshold for these smaller listed units than for subspecies or full species. Notably, all Pacific salmonids, which made up the majority of species involved in jeopardy determinations, are divided into multiple DPS/ESUs. It is also possible that NMFS, which manages fewer species than FWS, is able to dedicate more time and resources to each consultation, which in turn increases the agency’s confidence in finding and defending jeopardy determinations. Finally, differences between NMFS’s and FWS’s history and approach to consultations may explain some of our results^23^. Future research should evaluate the degree to which these factors are responsible for differences in how the ESA is implemented between the Services. While the causes of differences between the Services may be unclear, the low percentage of jeopardy and adverse modification findings shows that NMFS, like FWS, has worked with agencies and applicants to find solutions the vast majority of the time. Our results underscore the same message as research using parallel section 7 data from the FWS: “conventional wisdom” about the ESA stopping projects is unfounded^7^.

Administrative data are a key yet under-used resource for understanding the strengths and weaknesses of laws and policies and can be used to make their implementation more effective. To achieve better conservation outcomes for more species, conservationists have explored for decades more efficient approaches to administering the ESA. Data such as those that we evaluated here from NMFS provide critical insight into the reality of implementation and can inform regulatory policies (e.g., quantitative LAA guidelines) that improve conservation outcomes for imperiled species and make the ESA more efficient to implement. Despite its strength on paper, the ESA remains severely underfunded in practice^24^. As the number of endangered species continues to grow, and if Congress continues to cut funding for federal environmental programs, finding data-driven strategies becomes increasingly urgent. Especially in the face of funding shortcomings, the use of data rather than conjecture and anecdotes to guide policy and administrative decisions can provide options for consultation that efficiently and effectively conserve biodiversity. Our finding that proposed policies that minimize the involvement of expert agencies for the ESA could threaten the very existence of many listed species provides a stark illustration of the pitfalls of making policy decisions without data.

## METHODS AND MATERIALS

### Data Preparation

We obtained data from all formal and informal consultations as recorded in the PCTS database by NMFS biologists through June 2017. In addition to the species involved in a consultation and the determinations made by NMFS, PCTS records include the action agency, category of proposed action, dates of consultation initiation and conclusion, and the determinations proposed by action agencies. See supplemental Table S1 for a full list and description of fields.

Because records prior to 2000 were deemed potentially unreliable based on the frequency of data recorded and conversations with NMFS personnel, we analyzed data from 2000 to 2018. We performed several quality control steps to correct errors that may have accumulated from > 2,000 agency staff entering data over several decades. We corrected apparent date errors (e.g., end dates earlier than start dates) and homogenized the names of species, action agencies, and work types. NMFS records a variety of information about the nature of consultations in a single “Consultation Type” field. We split this into a “Type” field that indicates whether a consultation was recorded as formal, informal, or combined and a “Complexity” field that indicated whether a consultation was standard, programmatic, conference, or early.

Species and critical habitat determinations are recorded in a variety of combinations in PCTS. We standardized these outcomes by re-coding species determinations into one of four categories: ‘no effect’, ‘NLAA’, ‘no jeopardy’, or ‘jeopardy’. We re-coded critical habitat determinations into ‘no effect’, ‘NLAA’, ‘no adverse modification’, or ‘adverse modification’. We coded determinations for species that did not have critical habitat designated at the time of the consultation as ‘no critical habitat’. To ensure that all reported instances of jeopardy or adverse modification were accurate, we examined the biological opinions for these consultations and recorded proposed and final determinations, as well as work categories. Thus, our results reflect the minimum number of jeopardy determinations as there may have been erroneous non-jeopardy determinations recorded. In addition, we manually inspected 320 consultations for which outcomes were unclear based on PCTS records. Although this large dataset likely contains additional minor errors that we were unable to correct, we assume that those errors are unbiased and randomly distributed within the data.

### Data Analysis

All statistical analyses were performed in R 3.5.1^25^. We estimated changes in consultation frequency over time by fitting linear models with a log link and Poisson error distribution to the number of consultations recorded as informal and formal as a function of year. NMFS is organized into five geographic regions: Northeast, Southeast, Alaska, Pacific Island, and West Coast. We used a Chi-square test to estimate differences in formal consultation rates among geographic regions. Prior to 2013, the West Coast region consisted of the Southwest and Northwest regions, and we aggregated all consultations from these regions into a single West Coast category for consistency across years.

We also tested for differences in the frequency of consultation among species, action agencies, and work type using Chi-square goodness-of-fit tests, including only consultations for which a species was recorded. Out of 116 species consulted on by NMFS, 59 species had distinct population segments (DPSs) or evolutionarily significant units (ESUs) designated. For our analysis of species-specific consultation frequencies, we considered all DPS/ESUs of a given species together. For instance, a consultation involving multiple coho salmon (*Oncorhynchus kisutch*) DPSs was counted as a single consultation for coho salmon.

To evaluate patterns of agreement between NMFS and action agencies, we tabulated the frequencies of all possible combinations of determinations proposed by action agencies and those made by NMFS. We created an ordinal ‘discrepancy’ variable to rank the degree of disagreement between action agencies and NMFS. Determinations for which the action agency underestimated effects were assigned a negative score, while those in which the action agency overestimated effects were assigned a positive score (Table 1). Instances of agreement were assigned a score of ‘0’ and included situations in which both NMFS and the action agency determined no effect or NLAA, or when the action agency determined LAA and NMFS subsequently made either a jeopardy or no jeopardy determination (Table 1). We identified agencies exhibiting extreme rates of disagreement and agreement with NMFS using Kolmogorov-Smirnov tests comparing the distribution of discrepancy scores for a given agency against the distribution of discrepancy scores among all agencies. We restricted this analysis of discrepancy to agencies with at least 20 recorded consultations and considered an agency to exhibit significant departure from overall rates of disagreement if the probability of the test statistic *D* was *P* < 0.05.

We estimated the rate of jeopardy determination made during formal consultations, and the proportion of formal consultations with at least one jeopardy determination and tested for changes over time using generalized linear models. We tested for differences in jeopardy determination rates among species, and the proportion of consultations with a jeopardy conclusion among work types using Chi-square goodness-of-fit tests. We used matrix permutation to identify combinations of species and work categories that exhibited a disproportionately high frequency of jeopardy determinations. First, we constructed a matrix containing the frequency of jeopardy determinations for every combination of species and work type. To create a null distribution for these frequencies, we then randomized cell counts 1,000 times while keeping row and column totals fixed using the *vegan* package^*26*^ in R. The probability that an observed frequency was greater than random chance was calculated as the proportion of permutations in which the simulated frequency was greater than the observed frequency. We considered combinations with *P* < 0.05 to exhibit significant positive association, and report effect sizes as the difference between the observed and mean simulated cell frequencies.

Finally, a consultation determining jeopardy for one species may be more likely to also reach a jeopardy determination for other closely related and/or spatially proximate species. We quantified rates at which pairs of species were jeopardized by the same proposed action (i.e., in the same consultation), which we referred to as “co-jeopardization”. Rates of co-jeopardization that were greater or less than random were determined using a matrix permutation test. We organized consultation data into a binary species by consultation matrix in which cells indicated whether a species was jeopardized in a consultation. We estimated the pairwise probabilities of co-jeopardization and effect sizes (i.e., the difference between observed and expected frequency of co-jeopardization) for species with at least one jeopardy determination using the *cooccur* package^27^ for R. We considered pairs of species for which the proportion of permutations resulting in co-jeopardization was greater than observed *P* < 0.05 to exhibit significant association.

## ACKNOWLEDGEMENTS

We thank C. Tortorici and K. Peterson for providing access to the PCTS database. J. Miller, J. Rappaport Clark, and R. Dreher provided review and feedback on drafts of the manuscript.

